# Cross-regulation between proteome reallocation and metabolic flux redistribution governs bacterial growth transition kinetics

**DOI:** 10.1101/2023.07.06.547923

**Authors:** Huili Yuan, Yang Bai, Xuefei Li, Xiongfei Fu

## Abstract

Bacteria need to adjust their metabolism and protein synthesis simultaneously to adapt to changing nutrient conditions. It’s still a grand challenge to predict how cells coordinate such adaptation due to the cross-regulation between the metabolic fluxes and the protein synthesis. Here we developed a dynamic Constrained Allocation Flux Balance Analysis method (dCAFBA), which integrates flux-controlled proteome allocation and protein limited flux balance analysis. This framework can predict the redistribution dynamics of metabolic fluxes without requiring detailed enzyme parameters. We reveal that during nutrient up-shifts, the calculated metabolic fluxes change in agreement with experimental measurements of enzyme protein dynamics. During nutrient down-shifts, we uncover a switch of metabolic bottleneck from carbon uptake proteins to metabolic enzymes, which disrupts the coordination between metabolic flux and their enzyme abundance. Our method provides a quantitative framework to investigate cellular metabolism under varying environments and reveals insights into bacterial adaptation strategies.

## Introduction

Microbes are commonly exposed to changing conditions in their natural habitats [1]. To respond to fluctuating environments such as nutrient shifts, microbes redistribute their metabolic flux to synthesize amino acids and other internal metabolites, which are supplied as the precursors for the biosynthesis of protein. The variation in metabolic processes would further lead to the rebalance in allocation of protein synthesis among the catabolic proteins, ribosomal proteins, metabolic enzymes and others [2, 3]. On the other hand, the changes of protein allocation inversely alter the amino acids synthesis through the metabolic network [4]. Therefore, the cross-regulation between the cellular metabolic fluxes and proteome allocation under the changing environments provide a highly dynamical, intricate, and complex response.

Genome-scale metabolic network models (GEMs) are powerful tools for simulating cellular metabolism under different conditions. One of the most widely used methods for GEMs simulation is flux balance analysis (FBA), which optimizes a predefined objective function (such as biomass or metabolite production) subject to mass balance and capacity constraints [5]. FBA assumes that the fluxes including the nutrient uptake are at steady state, which limits its applications especially when the extracellular environment changes over time. Dynamic FBA (dFBA) extends FBA by incorporating the temporal variations of extracellular nutrients and metabolites, and simulate the metabolic adaptation of cells in response to environmental changes [6]. However, dFBA still neglects the effects of enzymatic constraints, which may be significant for energy metabolism [7, 8], metal utilization [9] and biomass productions [10].

Enzymatic constraints are essential for realistic simulation of metabolic models, but they depend on many parameters that vary over time and are hard to measure. Recent advances in proteomics data availability have enabled the integration of enzyme concentrations into the metabolic models, but other parameters, such as enzyme turnover rates or binding affinities, are still uncertain or unknown [10]. Moreover, these parameters are not constant, but change dynamically in response to environmental fluctuations. Therefore, the accuracy and applicability of such enzyme-constrained GEMs are limited by the availability and validity of these parameters. One way to overcome this challenge is to infer the dynamics of enzyme parameters from molecular interactions [11] and use them to predict the metabolic fluxes under changing conditions. However, this approach is computationally expensive and usually requires detailed knowledge of the gene regulatory network that controls enzyme expression.

Recent studies of quantitative bacterial physiology have revealed that bacterial gene expression is coordinated by functional sectors that respond to environmental changes [2, 12, 13]. The overall proteome can be coarse-grained into four major sectors based on their functions: carbon uptake (C-sector, *ϕ*_*C*_), metabolism (E-sector, *ϕ*_*E*_), translation (R-sector, *ϕ*_*R*_), and housekeeping (Q-sector, *ϕ*_*E*_) [14]. These sectors have different protein fractions that are regulated by the translational activity of the ribosome (*σ*), which in turn depends on the protein synthesis flux (*v*_*R*_) and the ribosome fraction. The global regulation of proteome resource allocation in steady-state growth follows the ‘growth law’ [8, 14, 15], while the flux-controlled regulation (FCR) can predict the kinetics of the protein reallocation without detailed metabolic fluxes under various carbon shift conditions [2].

Here, we developed a novel computational framework that integrates coarse-grained model of flux-controlled proteome allocation and genome-scale metabolic network model of flux balanced analysis to predict the metabolic flux redistribution during nutrient shifts. We applied our framework to analyze the temporal changes of individual metabolic fluxes of *Escherichia coli* during growth shifts between co-utilized carbon substrates. By comparing our predicted kinetics of metabolic fluxes with omics data of enzyme kinetics, we identified the bottleneck reactions among the metabolic network that limit the adaptation dynamics to nutrient changes. From a coarse-grained view, we found that the limiting factor for metabolic fluxes switched from carbon uptake proteins *ϕ*_*C*_ to metabolic enzymes *ϕ*_*E*_ during carbon down-shift. This switch resulted in a transient increase in the growth rate that was previously overlooked by experiments. Our results reveal the cross-regulation between proteome reallocation and metabolic flux redistribution plays an important role in the bacterial adaptation to environmental changes. Our framework provides a quantitative tool to investigate cellular metabolism under varying environments and reveals insights into bacterial adaptation strategies.

## Results

### Construction of the cross-regulated dynamical model

We developed a proteome-constrained metabolic model based on FCR and FBA to simulate the metabolic flux kinetics of *Escherichia coli* during carbon substrate shifts. The model was termed as dynamic constrained allocation flux balance analysis (dCAFBA). Specifically, we used the *E. coli i*JR904 genome-scale metabolic model [16] as the basis of the metabolic network, and introduced two coarse-grained constraints into the fluxes depending on their types of reactions (for example, *v*_*C*_ ≤ *ϕ*_*C*_/*w*_*C*_ for the carbon uptake flux (*v*_*C*_), and Σ_*i*_|*v*_*i*_| ≡*v*_*E*_ ≤ *ϕ*_*E*_ /*w*_*E*_ for the sum of all absolute metabolic fluxes (*v*_*i*_), where *w*_*C*_ and *w*_*E*_ were the protein efficiencies flux distribution *v*_*i*_ using the FBA by optimizing the biomass synthesis flux λ. We of the carbon uptake and metabolic proteins) (Fig. 1a). Therefore, we can obtain the then calculated the total amino acid synthesis flux (*v*_*aa*_) and considered it as the total protein synthesis flux (*v*_*R*_) under flux balance assumption. The total protein synthesis flux was further partitioned into the synthesis fluxes for the carbon uptake and translation according to FCR [2] (see Materials and methods).

**Fig. 1.**
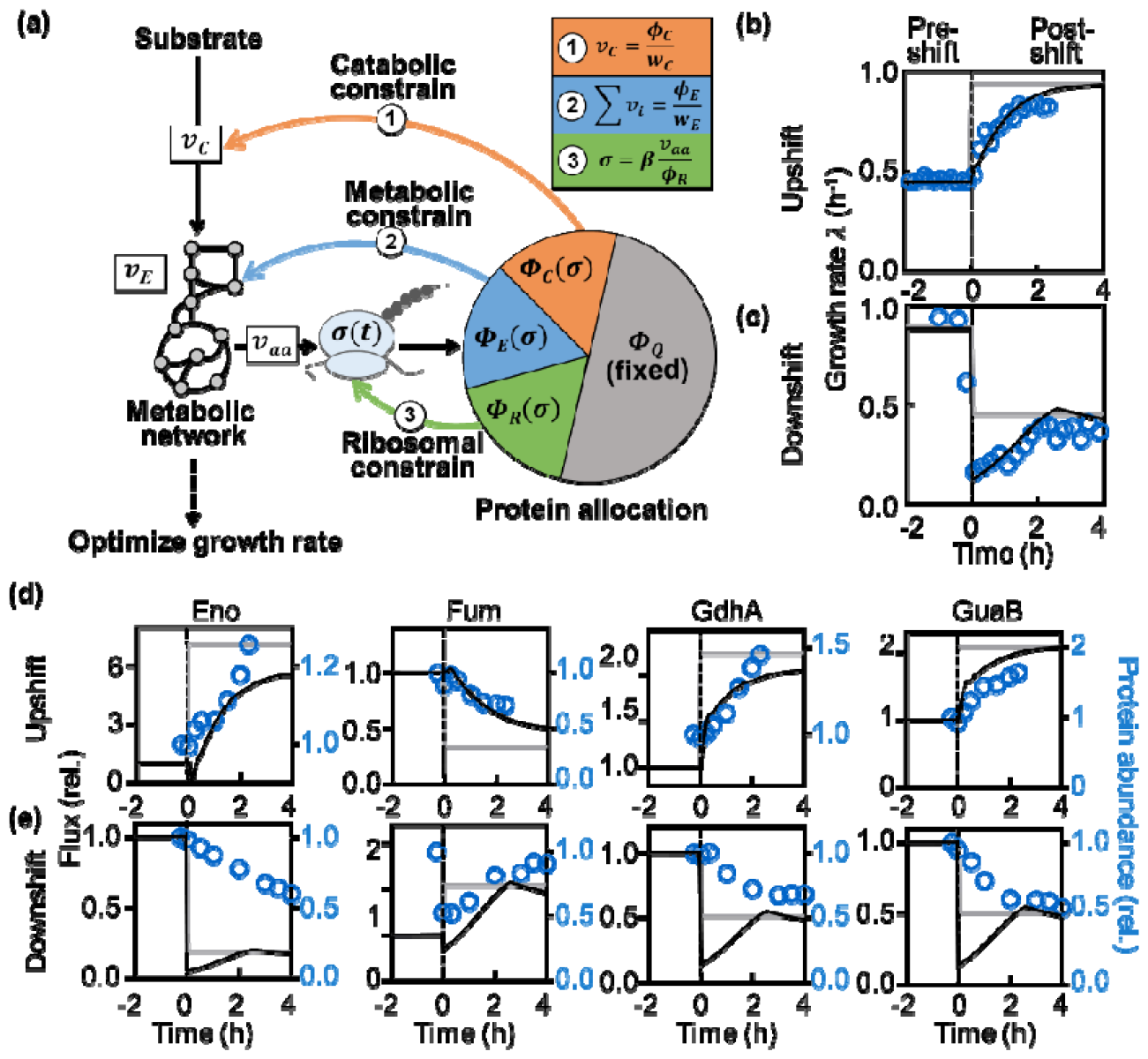
The dCAFBA model and its predictions during nutrient shift. (a) Schematic view of the model where the metabolic network (left) and the protein allocation are cross regulated. The carbon influx, constrained by protein partition of carbon uptake proteins, was converted to the synthesis flux of amino acids through the metabolic network which is constrain by the partition of enzyme proteins. The amino acids synthesis flux determines the translational activity of ribosome which then regulates the protein synthesis flux that are allocated to coarse-grained protein sectors such as carbon intake protein (C), metabolic protein (E), ribosomal protein (R) proteins and houses-keeping proteins (Q, fixed) via the regulatory function and (see Material and method). The dynamics of this model is assumed to be determined through, while the fluxes in the metabolic network are in quassi-steady state. The growth rate predicted by this model during up-shift of gluconate added to the medium of succinate (b, black line) and during down-shift of glucose depletion in the medium containing both succinate and glucose (c, black line) are compared to experimental data (blue circles, adopted from Erickson et al. [2]) and the predicted results of dFBA model (gray lines). The metabolic fluxes of typical reactions predicted by the dCAFBA model (black lines) during up-shift (d) and down-shift (e) are compared to the dynamics of related enzyme abundance (Eno for enolase; Fum for fumarase; GdhA for glutamate dehydrogenase; GuaB for Inosine-5’-monophosphate dehydrogenase) obtained from Erickson et al. [2] (blue circles) and the prediction of the dFBA model (gray lines).

To determine the metabolic fluxes during the nutrient shift, we adopted a quasi-equilibrium assumption that the metabolic fluxes reached a balanced state at each time step, as they adapt much faster than protein synthesis and growth mediated optimization and extracted the instantaneous amino acid synthesis flux *v*_*aa*_ (*t*)to dilution. We then obtained the dynamics of flux distribution by growth-rate further determine the protein reallocation kinetics (see Materials and methods). Therefore, this iterative framework enabled us to calculate the flux redistribution processes without the requirement of detailed knowledge about protein synthesis cost.

### Kinetics of metabolic fluxes during nutrient shifts

The dCAFBA model can first capture the transition kinetics of the growth rate and protein allocations during nutrient shifts. The growth rate is defined as the flux of biomass synthesis, which depends on the availability and utilization of nutrients. When bacterial population experience a growth up-shift (for example, gluconate added in medium of succinate in Fig. 1b, d & Fig. S1a, and another example in Fig. S2), they gradually increase their growth rate to a new steady state in a few hours (about 2-3 generations). In contrast, when they experience a down-shift (for example, glucose depleted from medium of glucose and succinate in Fig. 1c, e & Fig. S1b, and another example in Fig. S3), they quickly drop their growth rate and gradually recover to a lower post-shift state. During the transitions, cells change their gene expression patterns in response to changes in their growth conditions. Our dCAFBA model can also capture the relaxation kinetics of protein allocations, specifically in carbon uptake (*ϕ*_*C*_), translational (*ϕ*_*R*_) and metabolic (*ϕ*_*E*_) proteins (Fig. S1-S3). These transition kinetics are not easily captured by the classic dFBA model. dFBA model considers the nutrient availability as the limiting factor that governs the growth transition kinetics, thereby it only predicts immediately jumps of growth rate after nutrient shift (the grey lines in Fig. 1bc).

As an important improvement over the FCR model, our dCAFBA model can predict the transition kinetics of individual metabolic fluxes. Unlike the dFBA model, which suggests abrupt changes in the metabolic fluxes (the grey lines in Fig. 1b, c), our model revealed that the fluxes in the metabolic network would undergo a relaxation transition accompanied by the kinetics of enzymes during the carbon shifts. These detailed predictions of the transition kinetics of individual metabolic fluxes allows for closer examination of the transition kinetics of fluxes in the metabolic network, which further suggested that the transitions kinetics were not synchronized over the entire metabolic network.

To illustrate, we examined the reactions of the key pathways in the metabolic network, such as such as the reactions catalyzed by enolase (Eno) of the glycolysis pathway, fumarase (Fum) of the tricarboxylic acid cycle, glutamate dehydrogenase (GdhA) of the amino acid synthesis pathway, and insine-5’-monophosphate dehydrogenase (GuaB) of the nucleotide synthesis pathway. In both the up-shift and down-shift cases, the predicted transition time for these four reactions were consistent with the experimentally measured enzyme dynamics. In the up-shift case (Fig. 1d), the predicted fluxes of Eno, GdhA and GuaB increased gradually after a minor perturbation induced by the carbon shifts, while the flux of Fum decreases gradually. The metabolic fluxes of these reactions changed synchronously with the experimentally measured enzyme abundance related to these reactions [2]. In the down-shift case (Fig. 1e), the fluxes of all these reactions dropped abruptly after the carbon shift, and then increased smoothly until 2.4 hours and then decreased again to a steady value. The overall changes (from 0 to 4 hours) of the fluxes of the four reactions qualitatively coordinated with the changes of their related enzymes, where Eno, GdhA and GuaB decreased and Fum increased. However, the abrupt drop was not observed in the measured protein abundances, and the magnitude of the drop varied from one reaction to another. These complex behaviors highlight the need for a comprehensive understanding of the relationship between the dynamics of reaction flux and enzyme expression levels.

### Correlations between fluxes and enzymes

Based on the our model predictions, we further systematically examined the dynamic relationship between metabolic fluxes and their corresponding enzymes by calculating the temporal Pearson correlation coefficient (PCC) between each metabolic flux and its experimentally measured enzyme abundance after nutrient shift (see Materials and methods; Fig. S4). A positive PCC value indicates that the metabolic flux changes in the same direction as its reaction enzyme, while a negative PCC value indicates the opposite. Under different steady-state growth conditions, the majority of fluxes are positively correlated to the related enzyme abundance [17]. However, this relation is not always maintained especially during growth transition.

The temporal flux-enzyme correlation coefficients over the entire metabolic network revealed distinct patterns during the carbon up-shift (Fig. 2a) and down-shift (Fig. 2b). In the up-shift case, most reactions (Fig. 2c upper panel) exhibited positive flux-enzyme correlations, except for a few reactions related to the TCA cycle and ATP production pathways. The well-coordinated dynamics between metabolic fluxes and their enzyme abundance indicates that during carbon up-shift the carbon influx is surplus while enzymatic machinery limits the biomass synthesis fluxes, which requires both enzyme abundance and metabolic fluxes to gradually increase to adapt to their new steady state. In contrast, when cells underwent carbon down-shift, most reaction fluxes became negatively correlated with the changes of enzyme abundance (Fig. 2c lower panel). The negative correlation was a result of a gradual increase of fluxes after an abrupt drop at nutrient shift, while the related enzyme concentrations gradually decrease to the post-shift lower concentration state (Fig. 1e). This indicates that after down-shift the cells experienced a shortage of carbon influx which immediately results a drop in metabolic fluxes thereby the enzymes are surplus. These difference between up-shift and down-shifts further suggests that the efficiency of enzyme utilization would vary during the growth transitions.

**Fig. 2.**
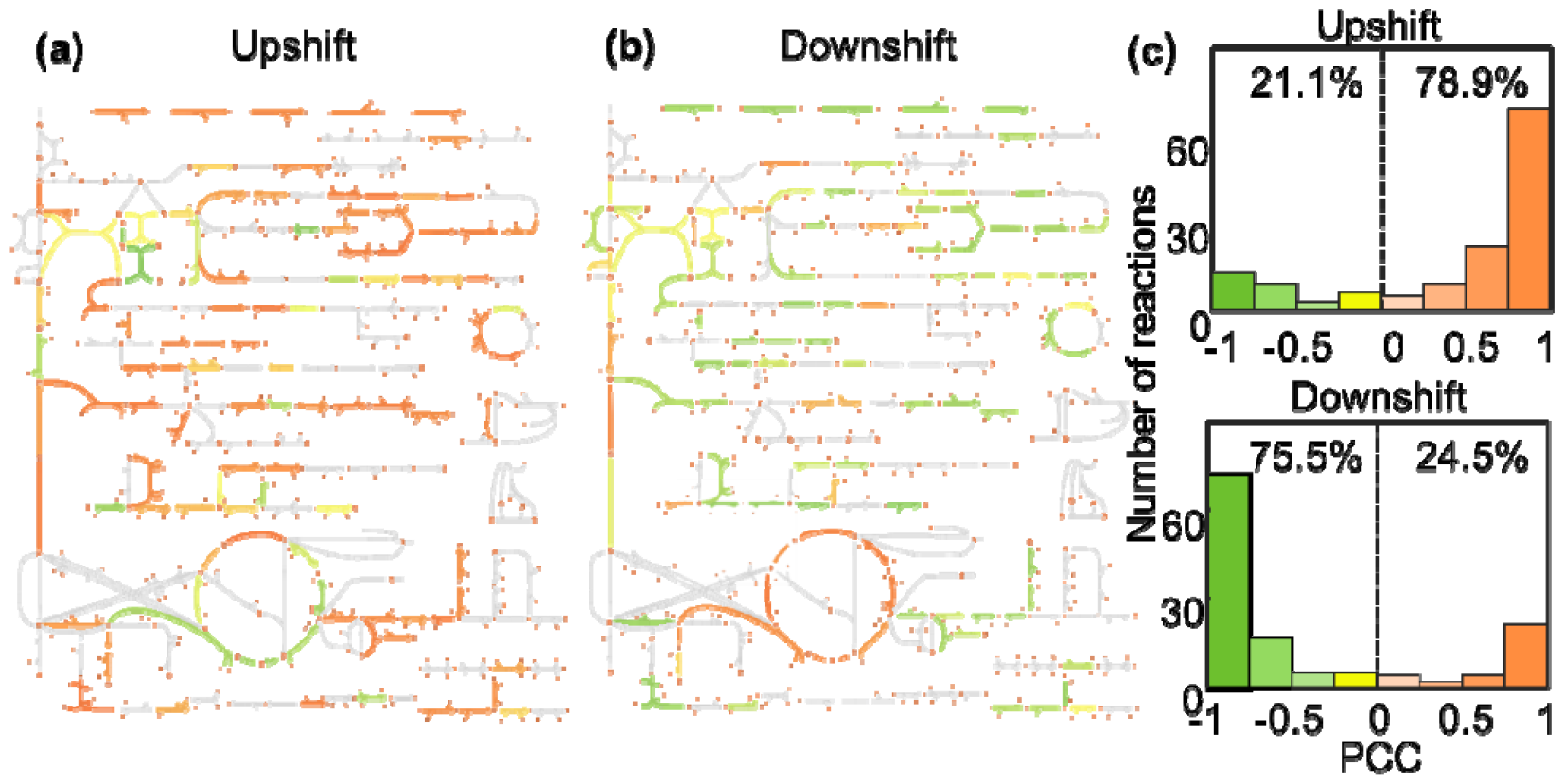
Illustration of the pearson correlation coefficient (PCC) between dynamics of the metabolic flux predicted by dCAFBA model and enzyme abundance during the up-shift case of gluconate added to the medium of succinate (a) and the down-shift case of glucose depletion in the medium containing both succinate and glucose (b). The calculation of PCC was presented in Materials and method. Colors from green to red represent the PCC value from -1 to 1. Gray color represents no data (nd). The histogram of the PCC values showed that 78.9% of the reactions have positive correlated dynamics between fluxes and enzyme abundance in the up-shift case, while only 24.5% of them are positively correlated in the down-shift cases (c).

### Bottleneck reactions during nutrient shifts

With the detailed prediction of the flux kinetics from the dCAFBA model, we are able to quantify the relative efficiency of enzyme utilization during the growth shifts, to determine the bottleneck reactions during nutrient shifts. The relative efficiency of enzyme utilization is defined as the ratio of the actual flux to the maximum possible flux given the enzyme abundance (see Materials and methods). During carbon up-shift, the reactions within glycolysis pathways exhibited a high level of efficiency in enzyme utilization, indicating that these enzymes play a limiting role in the kinetics of metabolic fluxes (Fig. S5). However, in the down-shift case, most of them showed low efficiency during the first 2-3 generation-time. The relative efficiency of enzyme utilization increased over time, as the fluxes of reactions adjusted to the post-shift steady state after the initial drops while the enzyme abundance decreased gradually (Fig. 3a). These results suggested the metabolic machinery is not always optimal during nutrient down-shift, due to the temporal constraints in nutrient influx.

**Fig. 3.**
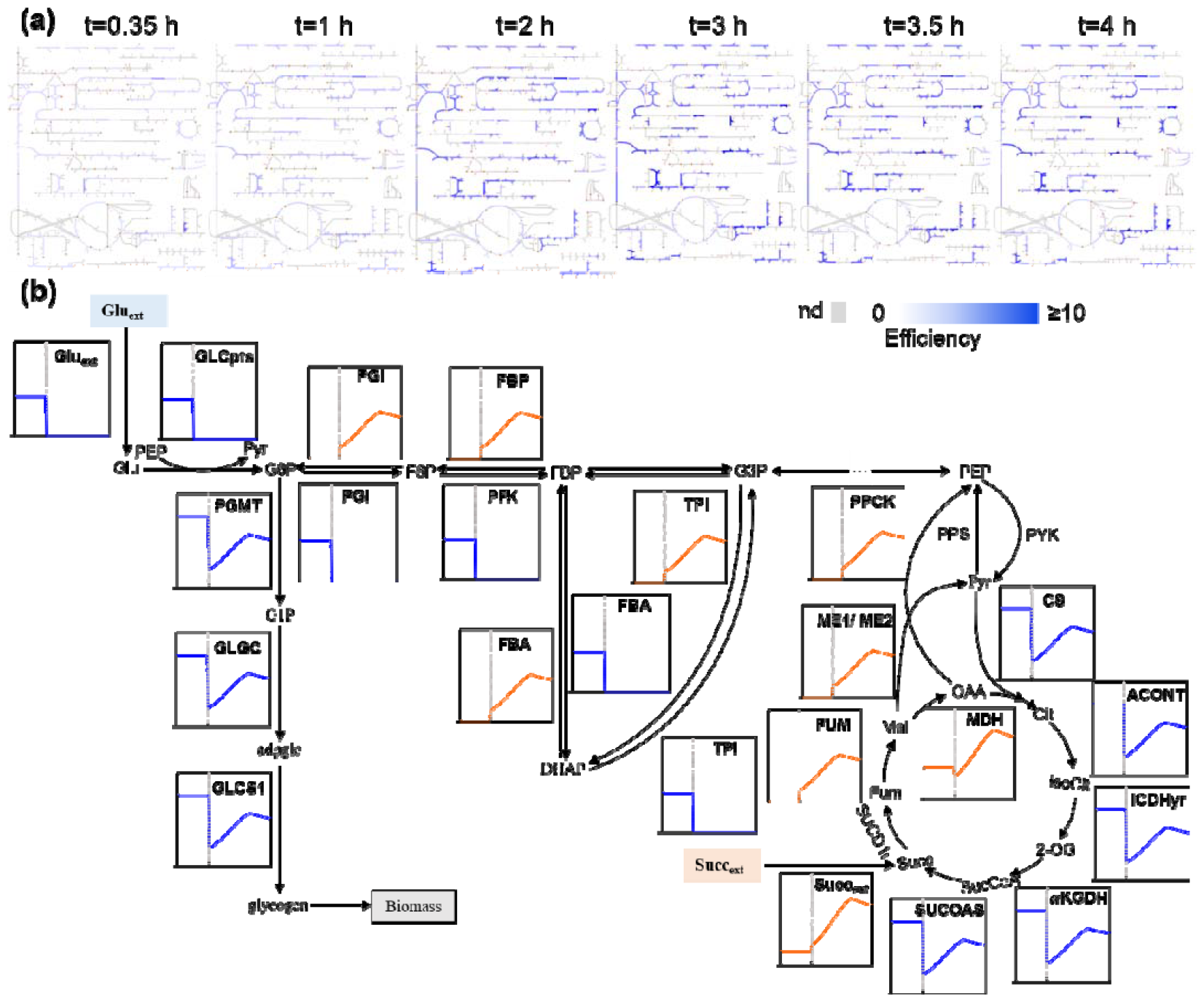
(a) Dynamics of protein efficiencies during carbon down-shift in the case of Fig. 1 (c). The deepness of the blue color represents the efficiency of enzyme utilization in the central metabolic network (see Materials and method for detailed calculation). (b) The dynamics of the selected reaction fluxes in the glycolysis, gluconeogenesis and TCA pathways were plotted. The x axis and y axis represent time (in hour) and reaction flux (mmol/gDW/h), respectively. The vertical grey line is the time at 0 hour, which separates the pre-shift (left) from the post-shift (right). The abbreviations for metabolites are as follows: Glu_ext_, extracellular glucose; Glu, glucose; G1P, glucose-1-phosphate; G6P, glucose-6-phosphate; F6P, fructose-6-phosphate; FBP, fructose-1,6-biphosphate; G3P, glyceraldehyde-3-Phosphate; DHAP, dihydroxyacetone phosphate; PEP, phosphoenol pyruvate; Pyr, pyruvate; OAA, oxalacetic acid; Cit, citrate; IsoCit, threo-isocitrate; Succ, succinate; Succ_ext_, extracellular succinate; SucCoA, succinyl-CoA; Fum, fumarate; Mal, malate; 2-OG, 2-oxoglutarate. The abbreviations for are as follows: GLCpts, glucose transporter; PGI, glucose-6-phosphate isomerase; PFK, phosphofructokinase; FBA, fructose bisphosphate aldolase; TPI, triose phosphate isomerase; PPS, phosphoenolpyruvate synthase; PYK, pyruvate kinase; CS, citrate synthase; ACONT, aconitase; ICDHyr, isocitrate dehydrogenase; AKGDH, 2-Oxogluterate dehydrogenase; SUCOAS, succinyl CoA synthetase; SUCD1i, succinate dehydrogenase; FUM, fumarase; MDH, malate dehydrogenase; PPCK, phosphoenolpyruvate carboxykinase; ME1/ME2, malic enzyme; PGMT, Phosphoglucomutase; GLGC, glucose-1-phosphate adenylyltransferase; GLCS1, glycogen synthase.

We further examined the temporal dynamics of the key catalytic reactions and their corresponding enzymes during the carbon down-shift of glucose depletion in medium of glucose and succinate. After glucose was rapidly depleted, the carbon influx switched to succinate uptake. The post-shift dynamics of succinate influx showed two phases (Fig. 3b): it first increased rapidly as the gene expression of succinate uptake proteins increased, and then decreased gradually to the new steady-state. The downstream reactions, such as succinate catalysis and malate synthesis (SUCC, SUCD and FUM), followed the same flux dynamic pattern. However, the fluxes of other reactions in the TCA cycle and biomass synthesis showed a different temporal pattern. They first dropped due to the lack of substrate supply, and then increased as the supply from upstream reactions gradually recovered. After 2-3 generation-time, those fluxes reached a peak and gradually adapted to the new steady state, as the limit factor shifted from succinate uptake to enzyme abundance in each reaction. This reduction in downstream fluxes fed back to upstream reactions, causing the decrease of succinate influx despite the continuous accumulation of its uptake proteins (Fig. S1). In summary, the flux pattern of the whole metabolic network during the down-shift can be divided into several sectors (for example, carbon intake and biosynthesis) depending on the limits of the reactions over time.

### Limitation switch during carbon down-shift

To address if the limitation switch depends on the specific reactions, we further proposed a coarse-grained picture of the metabolic network during the nutrient-shift (see Material and method). The changes of fluxes in the metabolic network can be categorized into two sectors, one that followed the dynamics of carbon intake flux *v*_*C*_, which was limited by the dynamics of the proteome allocation to the carbon uptake proteins *ϕ*_*C*_, and another that followed the overall metabolic flux defined by the sum of all the metabolic fluxes *v*_*E*_ ≡Σ|*v*_*i*_| and was limited by the dynamics of the proteome allocation to the metabolic enzymes *ϕ*_*E*_. Under assumption of the flux balance, the optimal flux distribution was constrained by the maximal fluxes which the two sectors could carry 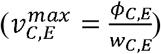.

In the up-shift case, the metabolic flux was limited by the maximal enzymatic flux 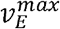 (Fig.S6), as the influx of high-quality carbon source increased the supply of substrates for central metabolism, while the downstream metabolic machinery was insufficient. Therefore, the cell increased the expression of metabolic enzymes 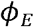, while the carbon uptake sector *ϕ*_*C*_ decreases. This resulted the decrease of 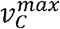and the increase of 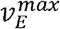, until a new balanced state was reached which allowed for an optimal growth.

During carbon down-shift, the cell experienced a bottleneck switch in the metabolic flux distribution (Fig. 4a & Fig. S7). At first, the glucose depletion caused an abrupt drop in both the fluxes and the maximal carbon influx, due to the limited pre-expressed uptake proteins for the new carbon source. Then, the carbon uptake protein sector *ϕ*_*C*_ gradually recovered, along with the increase of flux *v*_*C*_. After 2-3 generation time, the carbon influx was no longer the limiting factor, while the enzyme flux *v*_*E*_ reached its maximum flux 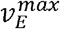 as the metabolic sector *ϕ*_*E*_ decreased due to the growth-mediated dilution. As a result, the maximum growth rate occurred at the time of limitation switch from carbon-uptake limitation to enzymatic limitation.

**Fig. 4.**
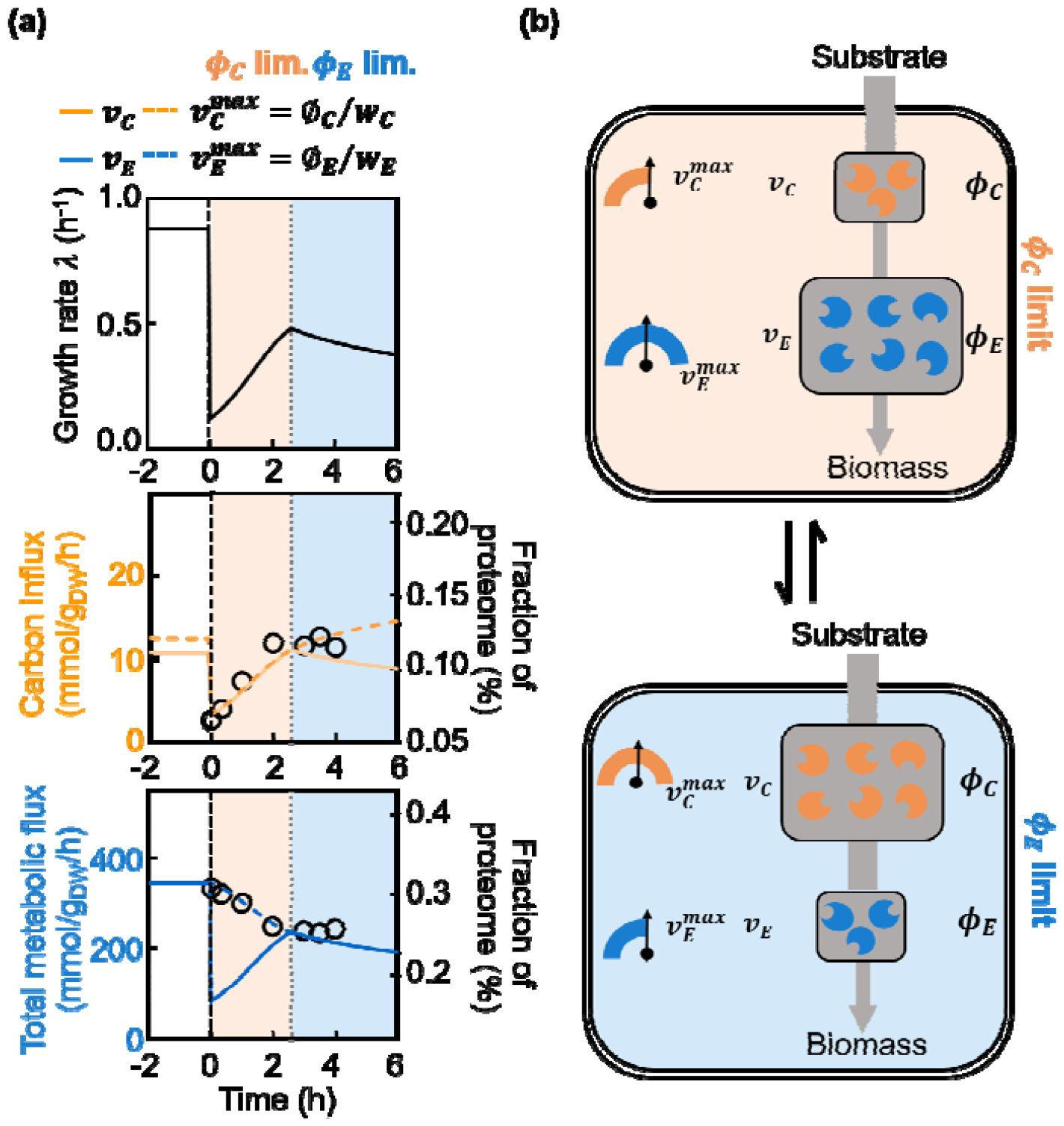
Limitation switch during carbon down-shift upon glucose depletion at t=0 in the medium containing both glucose and succinate. (a) The post-shift dynamics of the growth rate predicted by the dCAFBA model (upper panel) is divided into the carbon uptake protein limited region (limit, colored in orange) and the enzyme limited region (limit, colored in blue) at the peak of growth rate. These two regions have distinct dynamics relation between the actual carbon influx (solid orange line,) v.s. the maximal carbon influx (dashed orange line,) (middle panel), and the dynamic relation between the actual total metabolic flux (solid blue line,) and the maximum of metabolic fluxes (dashed blue line,) (lower panel). In this simulation, the following parameters were used: *w*_*c*1_= 1.96 x 10 gh/mmol, *w*_*E*_ = 8.3 x 10^−4^ gh/mmol and *h*_1_ =0.64, which allowed for the simulated half-life time for growth rate 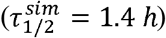was closely aligned to the experimentally measured one 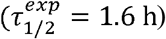. Where the half-life time was defined by the time-period to reach the middle value between pre-shift λ(*t=*0) and post-shift steady state λ_*f*_. (b) The *ϕ*_*C*_ limit and *ø*_*E*_ limit represent the fluxes of the coarse-grained reactions *v*_*C*_ and *v*_*E*_ is either limited by the maximal flux 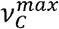or 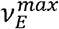 that can be carried by *ø*_*C*_ and *ø*_*c*_

Further analysis suggested that the time of limitation switch was determined by the protein efficiencies (). The lower uptake protein efficiency required less proteins to carry sufficient uptake flux to supply the metabolic system, leading to an earlier limitation switch. In contrast, higher efficiency of metabolic enzymes required more enzymes to carry the same metabolic fluxes in the pre-shift condition, resulting in a longer time for to decrease to match the recovery of carbon influx (Fig. S8). These results confirmed that the limitation switch during carbon down-shift was a result of global regulation of the protein sectors. The protein efficiencies, which determine the dynamic behaviors of the transition, would provide guidance for designing the growth transitions.

## Discussion

Bacterial cells respond to environmental perturbations by globally regulating their growth rate, gene expression and metabolism. Recent quantitative studies of bacterial physiology have provided a coarse-grained picture that capture the kinetics of proteome reallocation during nutrient shifts, yet it is still challenging to investigate the dynamics of metabolic network. Here, we combined flux balance analysis with the global regulation of proteome allocation to quantitatively predict flux redistribution processes. A coarse-grained approach was used to describe the proteome-remodeling kinetics while the optimization of the growth-rate was applied to determine the flux distribution. This dCAFBA method enabled us to predict the redistribution kinetics of metabolic fluxes by two free parameters (*w*_*C*_ and *w*_*E*_). This method allowed us to identify the bottleneck of the system during nutrient shifts. In the carbon up-shift cases, we showed that the accumulation of enzyme expression limited the dynamics of the flux redistributions, while the limiting factor switched from carbon uptake proteins to enzyme proteins during carbon down-shift.

We noted that our dCAFBA model predicted an overshoot of growth rate during carbon down-shift (Fig. 4a). This phenomenon has been seen in previous instances of carbon downshift but went unnoticed in experimental data (Fig. S9) [2]. This ubiquitous presence of this growth overshoot emphasizes the significance of the limitation switch during nutrient changes, which prior FCR models failed to capture. The study revealed that the limitation switch does not depend on the detailed knowledge of the metabolic network but on the unbalanced dynamics of *ϕ*_*C*_ and *ϕ*_*E*_ sectors. Hence, we proposed a modified FCR model (mFCR) to include the limitation switch by introducing the E sector proteins to constrain the total metabolic flux *ν*_*E*_ (see Materials and methods, Fig. S10a). Comparing the predictions from dCAFBA, mFCR and FCR models during carbon shifts, we observed that all the three models were capable to capture the growth dynamics during carbon up-shift (Fig. S10b upper panel). But only the dCAFBA and mFCR can reproduce the overshoot of growth rate during carbon down-shift (Fig. S10b lower panel).

In comparison with other existing models aiming to predict the kinetics of metabolic fluxes in fluctuating environments, such as the dFBA method [6], the deFBA model [18] and the dynamicME model [19], we concluded that our dCAFBA captured the kinetics much better than the dFBA model (Fig. 1 gray lines), while utilized much less unmeasured parameters of enzymes than the deFBA and dynamicME models. In addition to the *E*.*coli i*JR904 metabolic network [16], we also tested our dCAFBA model to latest metabolic network *i*ML1515 [20], yielding similar outcomes (Fig. S11). A list of related model parameters utilized for the simulations can be found in Table S4.

Quantitative understanding of the cross-regulatory kinetics between proteome reallocation and flux redistribution is a prerequisite for predicting cellular dynamics, such as the protein expression dynamics that are relevant to stress response [21], infectious diseases [22] and heterogeneous biosynthesis of valuable chemicals [23]. To address the latter, we constructed an extended dCAFBA model to simulate the cellular dynamics perturbed by the inducible exogenous genes expression for lycopene production in *E*.*coli* [24, 25]. To do so, we first included all the enzymatic reactions and transport reaction for lycopene production into the metabolic network and grouped them into a new lycopene production sector *ϕ*_*L*_. And then, we assumed that the lycopene synthesis flux *v*_*lyco*_was determined by the competition for the shared precursors between enzymes of *ϕ*_*L*_ and *ϕ*_*E*_ with 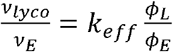. (Fig. S12a). the dynamics of *ϕ*_*L*_ was then determined by its synthesis flux χ_*L*_ in a similar way as *ϕ*_*R*_,*ϕ*_*C*_. Details of the model construction was presented in Materials and Methods.

Understanding how chemical production varies over time in response to different levels of exogenous gene expression following inducer addition is crucial when engineering external gene circuits. With this extended dCAFBA model, we can predict the kinetics of growth rate *λ*, all the reaction fluxes including the lycopene synthesis flux *v*_*lyco*_, and its protein fractions *ϕ*_*L*_ under induction of lycopene related gene expression (*χ*_*L*_ shifts from 0 to a preset value *χ*_*L*0_) (Fig. S12b). Our study demonstrated that the amount of lycopene produced within 24 hours of induction rises alongside pre-determined expression of lycopene-linked genes (*χ*_*L*0_) (Figure S12c). Nevertheless, the advantage derived from boosting gene expression declines as *χ*_*L*0_ increases. This is largely due to rapid decreases in growth rate under elevated *χ*_*L*0_ levels, constraining the increase of lycopene protein sector *ϕ*_*L*_. In summary, dCAFBA offers a useful blueprint for designing genetic circuits capable of adjusting key enzyme expressions in real-time to maximize metabolite yields [26-30].

## Acknowledgement

This work is partially supported by the National Key Research and Development Program of China (2018YFA0903400, 2023YFA0913900, 2021YFA0910703), the Strategic Priority Research Program of the Chinese Academy of Sciences (XDB0480000), NSFC (32071417), Guangdong Basic and Applied Basic Research Foundation (2022A1515110540), Shenzhen Science and Technology Program (ZDSYS20220606100606013).

## Author contributions

X.F. & Y.B. conceived the project. H.Y. performed all the numerical simulations. All the authors analyzed the results and wrote and revised the paper.

## Materials and methods

### Construction of the dCAFBA model

In this study, we performed all the model simulations in a proteome allocation model introduced by Mori et al. [31], which is constructed based on the *E*.*coli i*JR904 genome-scale metabolic model [16], including 761 metabolites and 1075 reactions. Following the recent studies on bacterial proteome [8, 14, 15], the entire cellular proteome was divided into four coarse-grained functional sectors in the proteome, which are carbon-scavenging proteins (C-sector) that includes proteins devoted to carbon import and transport from extracellular, metabolic enzymes (E-sector) that consists of biosynthetic enzymes, translational proteins (R-sector) that comprises ribosomal and its affiliated proteins, and house-keeping proteins (Q-sector) whose express keeps to be constant at various growth conditions. In the resulting proteome allocation model, there is a single reaction (carbon transport reaction) in C-sector. In total, 615 reactions belong to E-sector. 458 reactions were assigned to Q-sector. The fractions of the four proteome sectors should sum up to be 1, i.e. *ø*_*C*_+*ø*_*E*_+*ø*_*R*_+*ø*_*Q*_= 1.

The study of Erickson et al. [2] has shown that the protein synthesis flux of C sector and R sector are regulated by the translational activity *σ*(*t*)with regulation functions of *χ*_*C*_ (*σ*) and *χ*_*R*_ (*σ*)in the form of 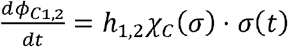 and 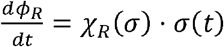, where the regulation functions of protein allocations *χ*_*C*_ (*σ*) and *χ*_*R*_ (*σ*) are: *χ*_*C*_ (*σ*)= (1−*σ*_χ*R*_*/λ*_*C*_*)ø*_*C,max*_, *χ*_*R*_ (*σ*) = ø_*R*,0_/(1−*σ*/γ).λ_*C*_ *γ,ϕ*_*C,max*_,*ϕ*_*R*,0_ are constants measured by the steady state protein allocationsunder different growth mediums [2] (see Table S2). The parameter *h*_1,2_ represents the partition of two co-utilized carbon sources used when both of them were presented in the growth medium. It is a fitted value that ensures the predicted growth rate equals the experimental data (Table S3). With the above regulation function and *χ*_*C*_, and *χ*_*R*_ the dynamics of each protein sector can be obtained: 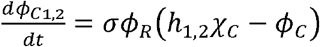 and 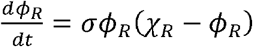. Under steady-state growth condition, the protein fraction of each sector *ϕ*_*C*1,2_, *ϕ*_*R*_ equals its respective regulation function *χ*_*C*_ and *χ*_*R*_. This is because the synthesis and dilution of each protein sector are balanced, which means 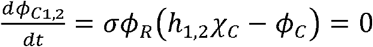 and 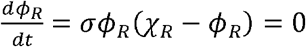. The dynamics of the metabolic enzymes *ϕ*_*E*_(*t*) can be written as the rest of the protein partition: *ϕ*_*E*_(*t*) =1−*ϕ*_*C*_−ϕ_*R*_(*t*) *;−ϕ*_*Q*_. With this knowledge, we can get the protein dynamics as long as the translational activity *σ*(*t*) is deduced.

Since the translational activity *σ*(*t*) ≡*v*_*R*_(*t*) / ϕ_*R*_(*t*) is defined by the protein synthesis rate (*v*_*R*_(*t*)) per ribosome (ϕ_*R*_(*t*)). We then assume all the synthesized amino acids from the metabolic network are assembled into protein synthesis, so that the protein synthesis rate is proportional to the amino acid synthesis rate *v*_*R*_(*t*) = *βv*_aa_(*t*) with a constant *β* to represent the conversion factor between moles and gram. As a result, the dynamics of *σ*(*t*) can be expressed as:

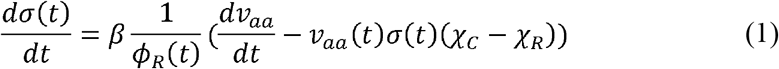

This equation describes how the amino acid synthesis flux *v*_*aa*_(*t*) regulates the dynamics of protein allocation *ϕ*_*C*_(*t*), *ϕ*_*R*_(*t*), *ϕ*_*E*_(*t*) via translational activity *σ*(*t*). It is also important to formulize how protein allocation constrains the metabolic flux distribution through the metabolic network. To do so, we simply set the upper boundaries on the carbon uptake flux *vC* and the sum of all the absolute metabolic fluxes *Σ*_*i*_ |*v*_*i*_| by:

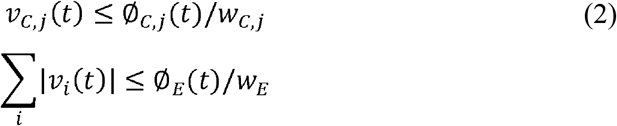

where *i,j* are the index of metabolic reactions in E-sector and index of carbon exchange reactions, respectively. *w*_*C,j*_ and *w*_*E*_ characterizes the carbon uptake protein and metabolic enzymes needed to carry one unit flux, which are determined by fitting the growth rates at steady state and protein partitions at corresponding growth medium (Table S3). With protein allocation constrains, we can calculate the flux distribution *v*_*i*_(*t*) by optimizing the biomass synthesis flux *λ*.

Based on experimental findings [15], *Ø*_*C*_ was assumed to depend linearly on the carbon intake flux *v*_*C*_, i.e. Ø_*C =*_ Ø_*C*,0_ + *w*_*C*_ *v*_*C*_, where Ø_*C*,o_ represents a growth rate-independent offset and is assumed to be 0 in this work, unless otherwise stated elsewhere. *w*_*C*_ characterizes the efficiency of the carbon catabolic protein for carbon intake. *v*_*C*_ is the carbon influx that carbon transport reaction carries.

Similarly, the proteome fraction of the E-sector is written as Ø_*E*_ = Ø_*E*_._0_ + *w*_*E*_Σ_*i*_|*v*_*i*_|, where Ø_*E*,o_ is the growth rate-independent offset and is also assumed to be a 0 in this work. The weight *w*_*E*_ stands for the efficiency of a metabolic enzyme for carrying one unit of metabolic flux *v*_*i*_. In this study, *Ø*_*Q*_ of 0.45 reported in a previous work [14] has been exploited. Other parameters for simulating the cell growth during the shift are listed in Table S2.

### Model simulations

All simulation was performed using FBA as implemented in the COBRA toolbox executable in MATLAB. The GLPK Optimizer solver in combination with the toolbox was used to solve the linear programming problems. In this study, the biomass equation was used to obtain the optimal solution of the metabolic model as described elsewhere [5]. Mathematically, the dCAFBA can be represented as follows:

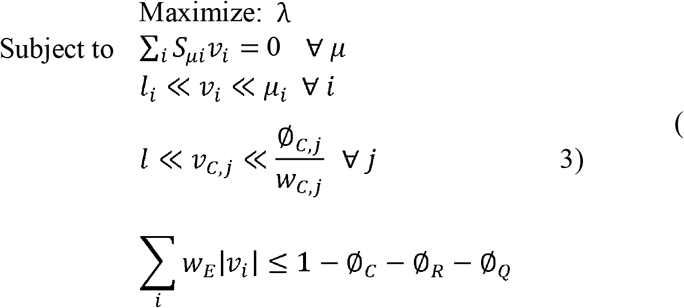

where λ is the flux that the biomass reaction carries, representing the growth rate, *S*_*ui*_ is the stoichiometric matrix, is a vector of all metabolites (u) and reactions (i), *l*i and *u*i represent lower and upper bounds for the flux of each reaction *v*_*i*_, respectively. *V*_*c*_ is the carbon influx that carbon transport reaction carries, Ø_*C*_ represents proteins associated with carbon intake and transport. The constants of the substrates used in this work are given in Table S3.

### Calculation of Pearson correlation coefficient

In this manuscript, we used the Pearson correlation coefficient to quantify the similarity between the dynamics of metabolic fluxes *v*_*i*_(*t*) and their corresponding enzymes *ϕ*_*i*_(*t*) after carbon shift (*t* >0). The calculation was performed using a Matlab command corr which calculated the statistical measure of the strength and directionality of the linear relationship between two continuous variables *v*_*i*_(*t*) and *ϕ*_*i*_(*t*) with formula of 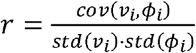 where *cov* (*v*_*i*_, *ϕ*_*i*_) represents the covariance while *std*(*v*_*i*_) and *std*(*ϕ*_*i*_) represent the standard deviations of *v*_*i*_ and *ϕ*_*i*_. The Pearson correlation coefficient “*r*” falls between -1 and +1, where positive values indicate a positive relationship between fluxes and enzymes (as one factor grows over time, so does the other).

### Efficiency of enzyme utilization

The relative efficiency *ϵ*_*i*_(*t*) of the enzyme utilization was defined by the ratio between fluxes *v*_*i*_(*t*) and the enzyme partitions *ϕ*_*i*_(*t*) of each reaction *i* normalized by the pre-shift enzyme efficiency 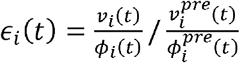 where 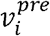 and 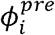 denotes the fluxes and enzyme partitions in the pre-shift steady state. In our calculation, the fluxes of a given reaction was predicted by the dCAFBA model while the protein partition of the enzymes are measured by Erickson et al. [2].

### Construction of the simplified model

The modified FCR model is built on the published FCR model [2] by introducing the dynamics of E protein: *ϕ*_*E*_(*t*)=1 −*ϕ*_*R*_(*t*)−*ϕ*_*C*_(*t*)−*ϕ*_*Q*_ shown in Figure 4b. We assume the protein synthesis flux *v*_*R*_ to be the amino acid synthesis flux *v*_*aa*_, the translational activity *σ* in the context can be described as 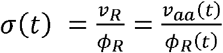.

According to the results of our dCAFBA model, we set constraints on the amino acids synthesis flux *v*_*aa*_ either by the amount of carbon uptake proteins *ϕ*_*C*_ or the amount of metabolic proteins *ϕ*_*E*_ with a simple piece-wise function shown in equation (3).

When μ_*f*_, *ϕ*_*C*2_ (*t*)≤α*ϕ*_*E*_ (*t*), the differential equation for *σ*can be expressed as 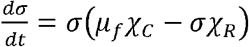. While, if μ_*f*_, *ϕ*_*C*2_ (*t*)≤α*ϕ*_*E*_ (*t*), and considering that *σϕ*_*R*_ =α*ϕ*_*E*_ =α(1 −*ϕ*_*R*_*−ϕ*_*C*_*−ϕ*_*Q*_), the differential equation for *σ* is written as 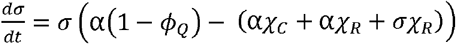 The constants α appearing in the modified coarse-grained model determined by fitting to the growth rate at steady state. Besides, the initial growth rate λ_*I*_ and the final growth rate λ_*f*_ were required in the modified FCR model, to are determine initial condition of the translational activity 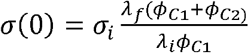 and the maximal carbon uptake rate 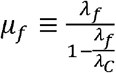 in the post-shit medium with λ_*C*_*=*1.17*h*^*−1*^

### Construction of the extended dCAFBA

To study the cellular dynamics perturbed by the inducible exogenous gene expression, we developed an extended model on the basis of *E*.*coli i*JR904 [16]. We added three enzymatic reactions (IPDPTT, GGDPT, FADO) from Choi et al. [32] and transport reaction (LYCO, transporting lycopene from cytoplasm to environment) into the metabolic network *i*JR904. These newly added reactions were assigned to a new *L* sector with the related to lycopene synthesis proteins were defined as *ϕ*_*L*_. This new metabolic network, termed Model_iJR904Lyco, is available in the simulation code. Considering that there is an additional flux drain included in the metabolic model, the exogenous and endogenous enzymes would compete for the shared precursors such as glyceraldehyde 3-phosphate (G3P) and pyruvate. We assume that the fluxes going into exogenous and endogenous metabolic reactions are allocated linearly with the ratio of their relative protein sectors, i.e. 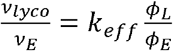. The parameter *k*_*eff*_ represents the competition for the shared precursors between lycopene synthesis and other metabolic pathways. The illustration of the expanded model was shown in Fig. S12 (a). The model parameters used in the simulation is given in the Table S5.

In our extended model, a step function χ_*L*_ which controls the lycopene protein synthesis was introduced 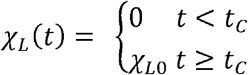,to present the induction of lycopene synthesis gene expression at time *t*_*C*_ to level χ_*L*O_. Similar to the *ϕ*_*C*_ (*t*)and *ϕ*_*R*_(*t*), the protein fraction related to the lycopene synthesis is also determined by the translational activity *σ*(*t*) in the form of 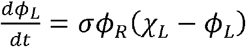. Once the and *ϕ*_*L*_ (*t*_*n*_) and *ϕ*_*E*_ (*t*_*n*_) at each time step *t*_*n*_ have been determined, the lycopene synthesis flux *v*_*lyco*_ (*t*_*n*_) can be calculated using the total metabolic flux from the previous time step *v*_*E*_ (*t*_*n−1*_). To determine the dynamic metabolic fluxes, we then solve FBA system with an optimization of growth rate λ (*t*_*n*_)subject to the lycopene synthesis flux *v*_*lyco*_ (*t*_*n*_).

## Data availability

Data supporting the findings of this study are available within the main text and Supplementary Information. The Custom-made simulation code is available at github: https://github.com/BaiYangBqdq/dCAFBA

## Supplemental tables

**Table S1:** Description of variables.

| symbol | description |
| --- | --- |
| $\sigma$ | translational activity |
| $\chi_{Rb}$ | regulation function of ribosomal protein allocation |
| $\chi_C$ | regulation function of catabolic protein allocation |
| $\phi_{Rb}$ | fraction of ribosomal protein over total protein |
| $\phi_C$ | fraction of catabolic protein over total protein |
| $v_{aa}$ | protein synthesis flux |
| $v_C$ | carbon uptake rate |
| $\lambda$ | cell growth rate |

**Table S2:** Description of parameters. Default values were used unless otherwise stated.

| symbol | description | value | source |
| --- | --- | --- | --- |
| $\gamma$ | Inverse slope of growth laws | 13.52 h<sup>-1</sup> | [2] |
| $\lambda_C$ | Strain specific constant of growth laws | 1.17 h<sup>-1</sup> | [2] |
| $\phi_{Rb,0}$ | A growth rate-independent offset of R sector | 0.049 | [2] |
| $\phi_Q$ | Fraction of house-keeping protein | 0.45 | [14] |
| $\phi_{C,max}$ | Maximum of proteome attainable to C, E and R sectors | 0.44 | [15] |
| $\beta$ | conversion factor between moles and gram | 0.1968 g/mmol | |
| $h_{1,2}$ | carbon-specific allocation | | |
| $\omega_E$ | enzyme cost per unit metabolic influx | 8.3×10<sup>-4</sup> gh/mmol | [31] |
| $\omega_C$ | enzyme cost per unit carbon influx | See Table S3 | |

**Table S3:**
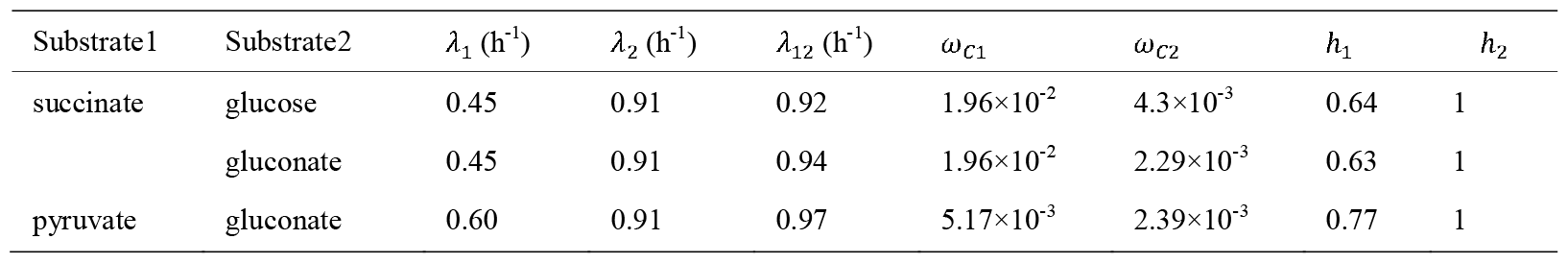
The constants of the substrates used in the simulation of dCAFBA.

**Table S4:** The parameters used in the dCAFBA simulations with the *E*.*coli i*ML1515 metabolic model.

| | Substrate1 | Substrate2 | $\omega_{C1}$ (gh/mmol) | $\omega_{C2}$ (gh/mmol) | $\omega_E$ (gh/mmol) | $\beta$ (g/mmol) | $h_1$ | $h_2$ |
| --- | --- | --- | --- | --- | --- | --- | --- | --- |
| Up-shift | succinate | gluconate | 1.96×10<sup>-2</sup> | 2.1×10<sup>-3</sup> | 3.3×10<sup>-4</sup> | 0.1870 | 0.63 | 1 |
| Down-shift | succinate | glucose | 1.6×10<sup>-2</sup> | 2.9×10<sup>-3</sup> | 5.15×10<sup>-4</sup> | 0.1870 | 0.6 | 1 |

**Table S5:** The parameters used in the simulations of lycopene production by dCAFBA.

| Substrate | $\lambda_1$ (h<sup>-1</sup>) | $\omega_C$(gh/mmol) | $\omega_E$(gh/mmol) | $\phi_{Cmax}$ | $k_{eff}$ | $t_C$ (h) |
| --- | --- | --- | --- | --- | --- | --- |
| glucose | 0.85 | $1.0\times 10^{-3}$ | $4.1\times 10^{-4}$ | 0.24 | 0.05 | 0.6 |

## Supplemental figures

**Fig. S1.**
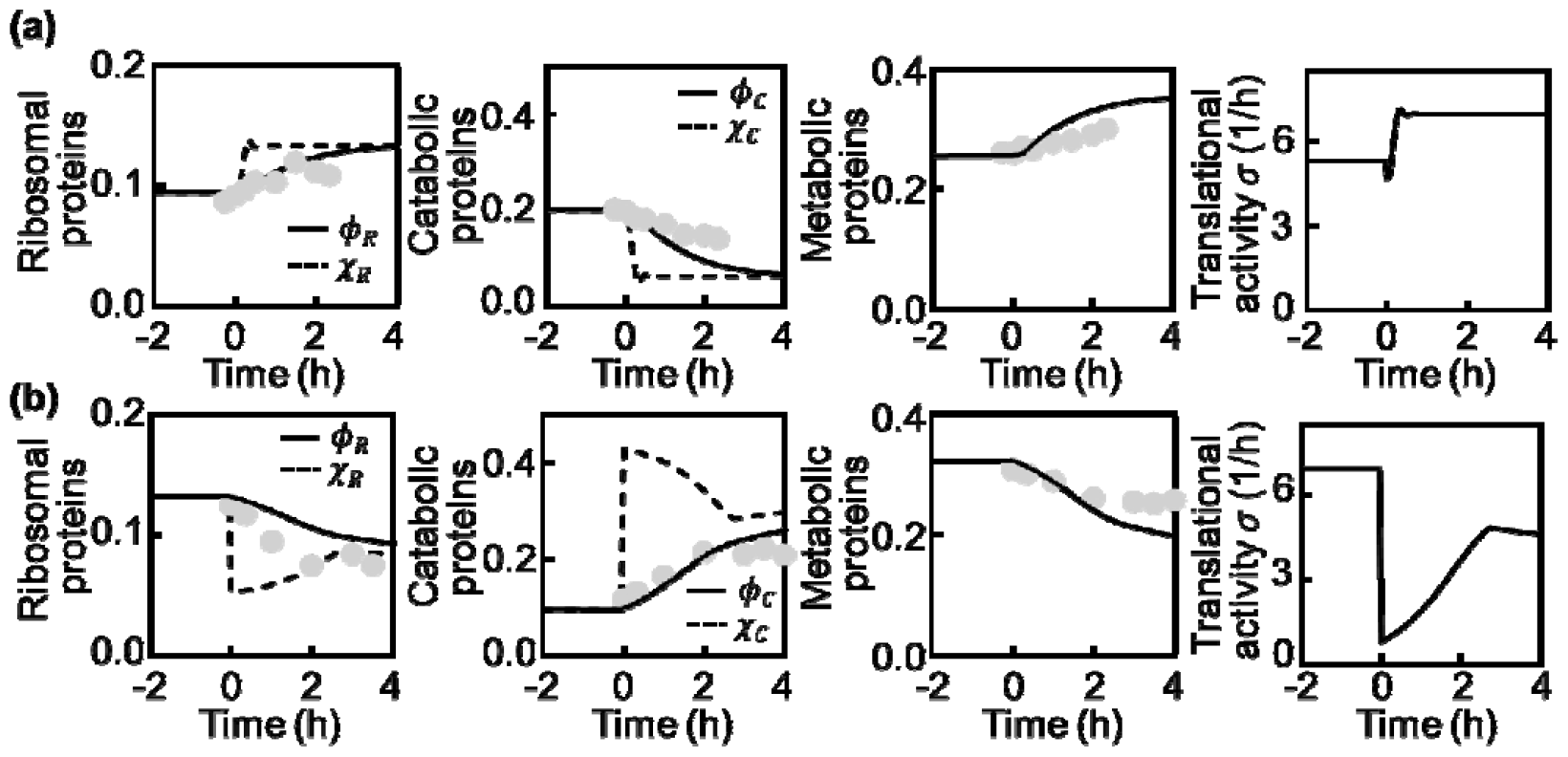
Regulatory functions and protein fractions of catabolic proteins, ribosomes, metabolic proteins and translational activity during the up-shift (a) and down-shift (b) of Fig.1. Dots, experimental data obtained from Erickson et al. [2]. Black lines, simulations of dCAFBA.

**Fig. S2.**
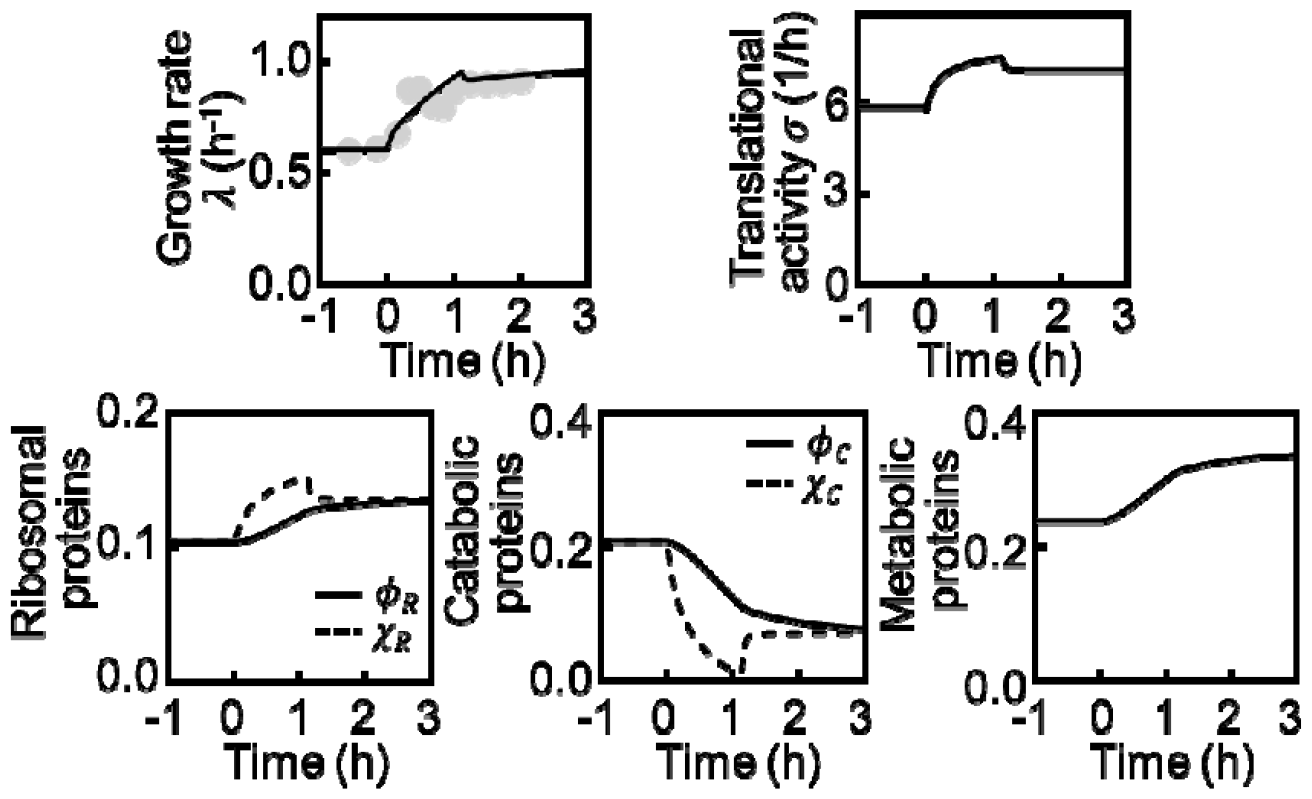
Growth rate, translational activity, and regulatory functions and protein fractions of catabolic proteins, ribosomes as well as metabolic proteins during the up-shift of gluconate added to the medium of pyruvate. Dots, experimental data obtained from Erickson et al. [2], if available. Black lines, simulations of dCAFBA.

**Fig. S3.**
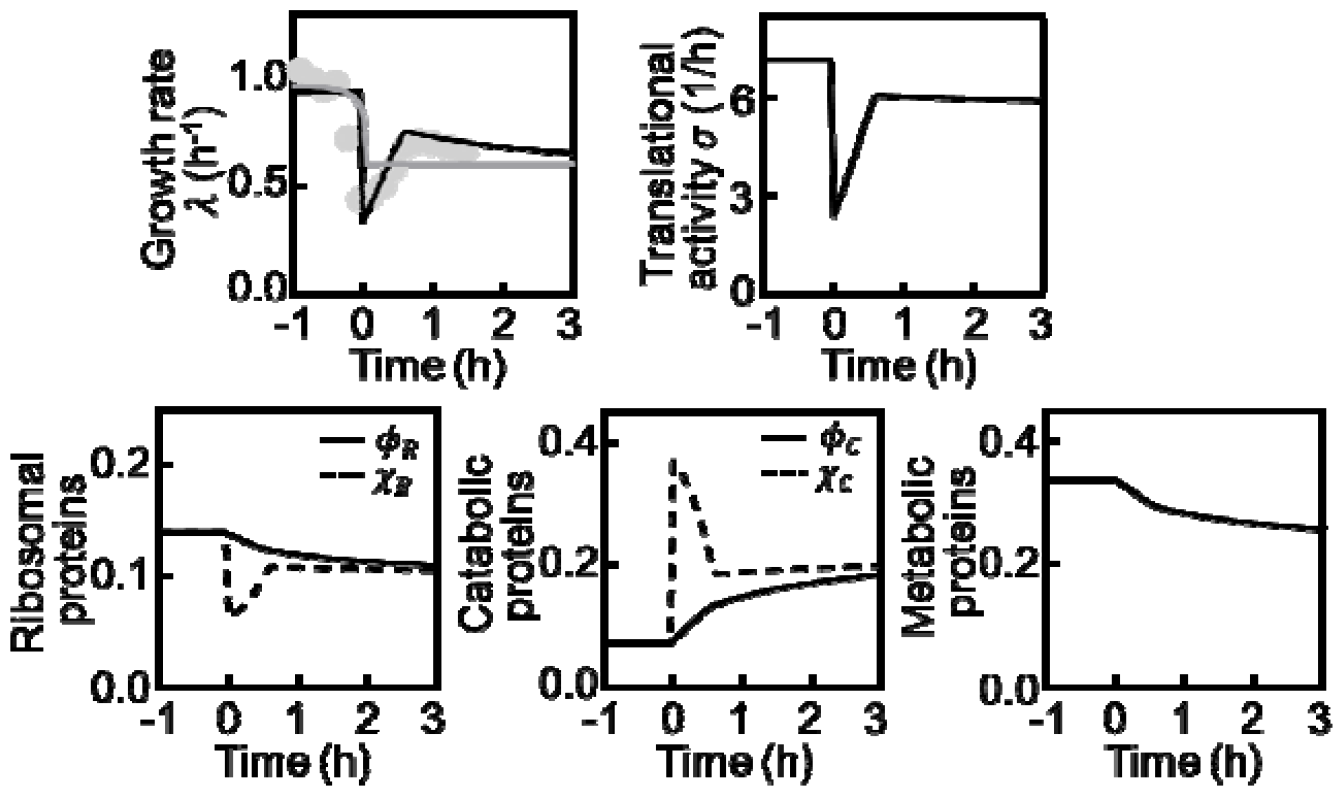
Growth rate, translational activity, and regulatory functions and protein fractions of catabolic proteins, ribosomes as well as metabolic proteins during the down-shift of gluconate depletion in medium of gluconate and pyruvate. Dots, experimental data obtained from Erickson et al. [2], if available. Black lines, simulations of dCAFBA. Grey line in the upper left panel represents the prediction of dFBA.

**Fig. S4.**
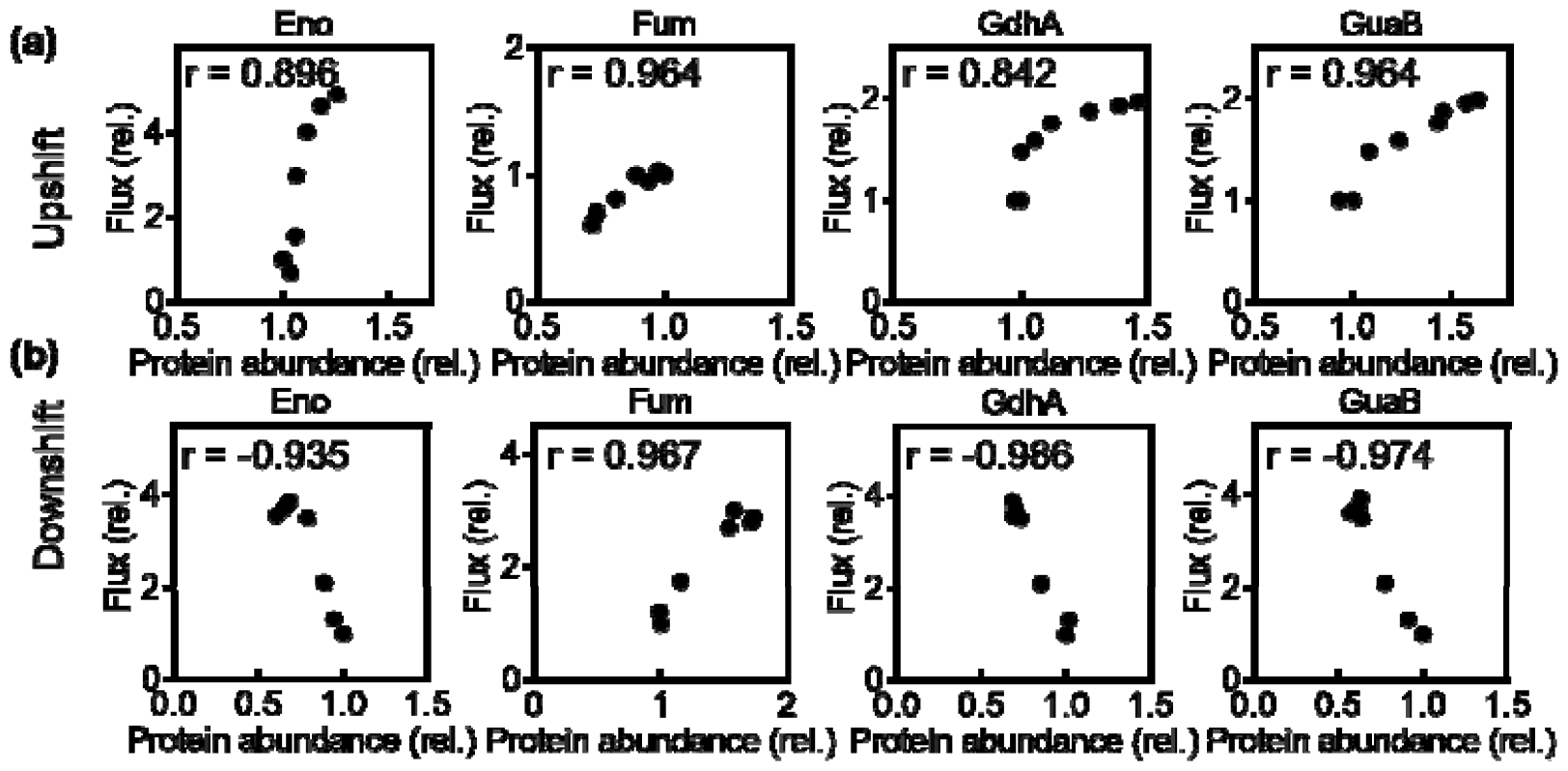
Correlations between predicted metabolic fluxes and experimentally measured enzyme abundances of selected reactions during up-shift (a) and down-shift (b). Each dot represents the flux and enzyme abundance at a time point. Symbo represents the Pearson correlation coefficient. Eno for enolase; Fum, for fumarase; GdhA for glutamate dehydrogenase; GuaB for Inosine-5’-monophosphate dehydrogenase.

**Fig. S5.**
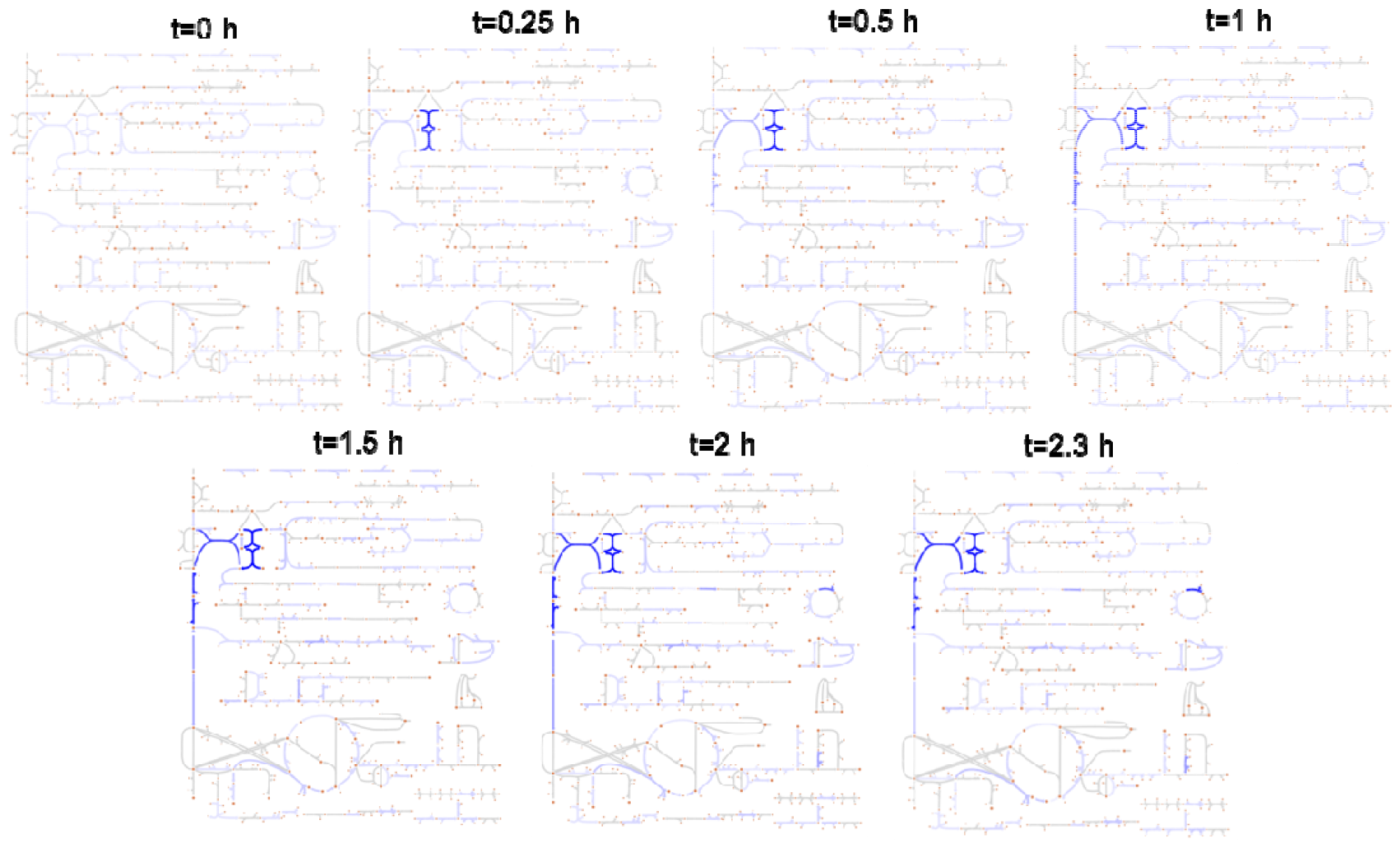
Dynamics of the protein efficiencies during carbon up-shift case of gluconate added to the medium of succinate as in Fig.1b. The deepness of the blue color represents the value of protein efficiency calculated by, where is the predicted flux of a given reaction in the metabolic network and is the protein partition of the respective enzyme measured by Erickson et al.[2].

**Fig. S6.**
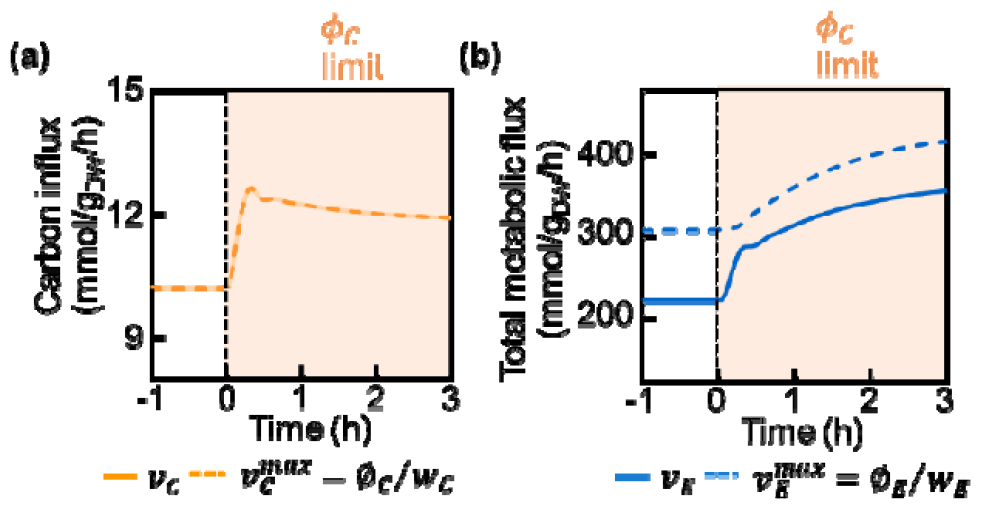
Comparisons between actual flux and maximal fluxes in the up-shift case of gluconate added to the medium of succinate as in Fig.1b. (a) Carbon influx (solid orange line,) v.s. the maximal flux the carbon uptake protein can carry (dashed orange line,). (b) Total metabolic flux (solid blue line,) v.s. the maximal metabolic flux the enzyme proteins can carry (dashed blue line,). These showed that the post shift period dynamics was limited by carbon intake proteins.

**Fig. S7.**
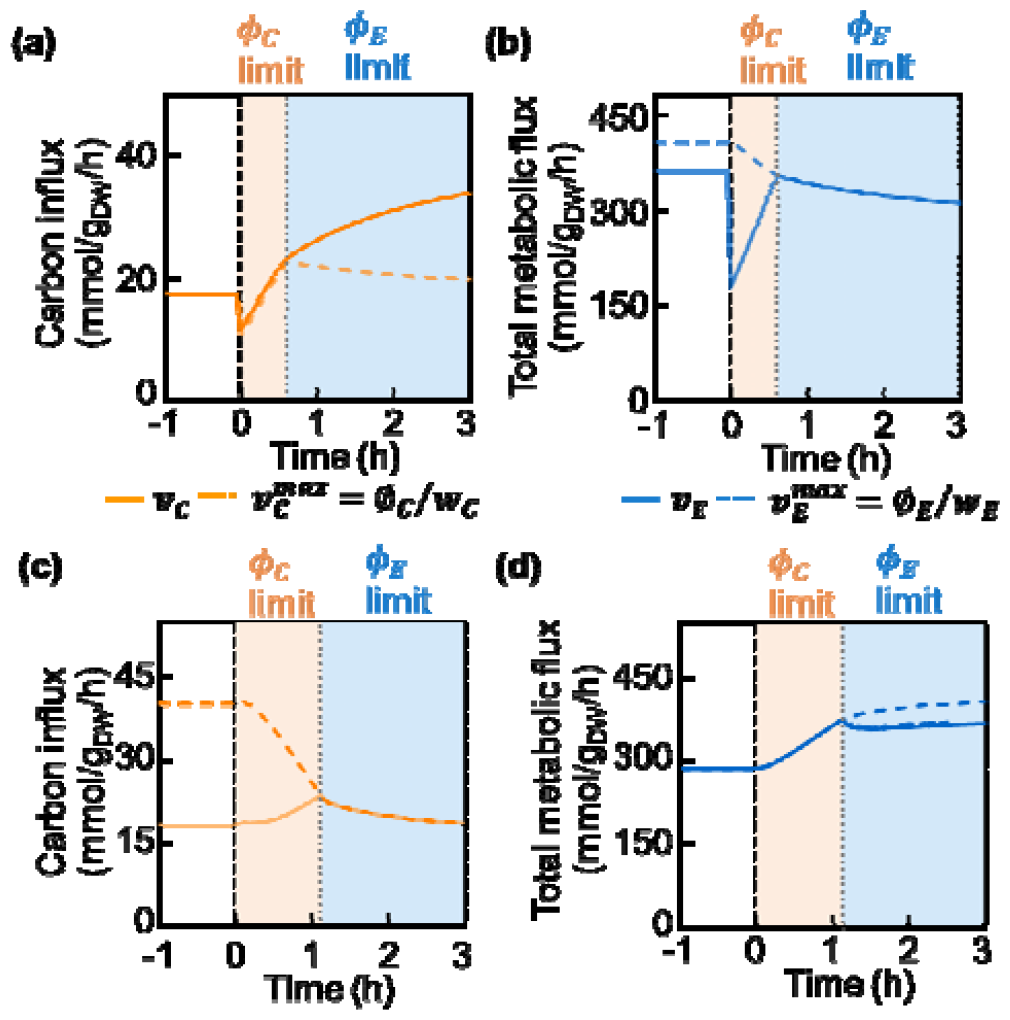
Comparisons between the actual flux (solid lines) and the maximal flux determined by enzyme abundance (dashed lines), for carbon intake flux (orange lines) and the total metabolic flux (blue lines), in the case of down-shift upon depletion of gluconate in the medium containing both gluconate and pyruvate (a, b) and up-shift upon the addition of gluconate to the medium containing pyruvate (c, d). Orange and blue regions represent the growth period was limited either by the carbon intake proteins (limit) or the enzyme proteins (limit), respectively.

**Fig. S8.**
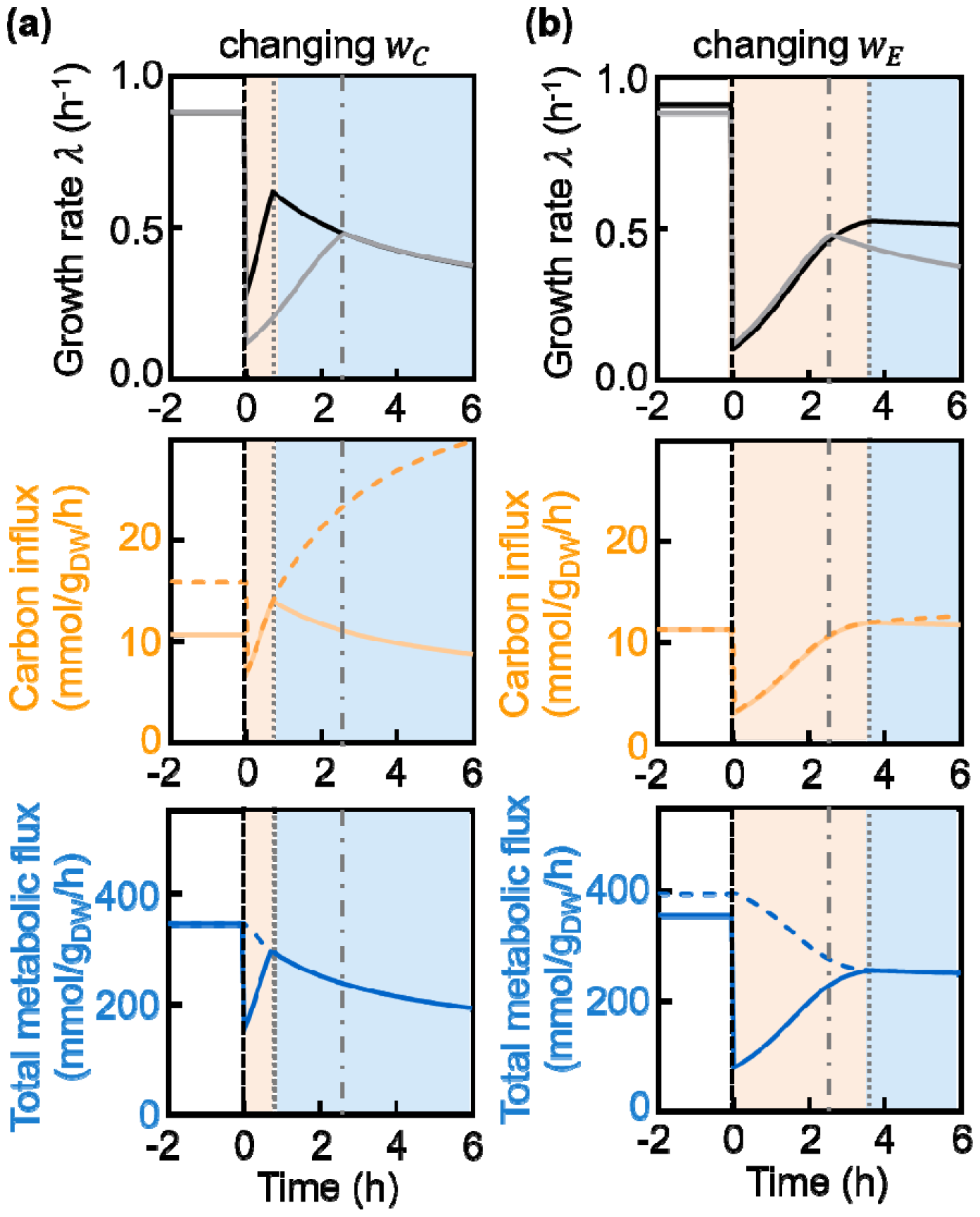
Effects of parameters (a) and (b) on the model predictions in the down-shift case upon depletion of glucose (depleted around at t=0), succinate (present continuously). The gray lines are the predicted results using the parameters as in Fig. 4a. Solid orange line is the carbon influx ; dashed orange line is the maximum of carbon influx, ; solid blue line is total metabolic flux ; dashed blue line is the maximum of metabolic fluxes. Gray dot-dashed lines in all panels indicate the time of limitation shift. = 0.96 10^−2^ gh/mmol, = 8.3 10^−4^ gh/mmol and =0.64 are used in the simulation of (a). = 1.96 10^−2^ gh/mmol, = 9.5 10^−4^ gh/mmol and =0.64 are used in the simulation of (b).

**Fig. S9.**
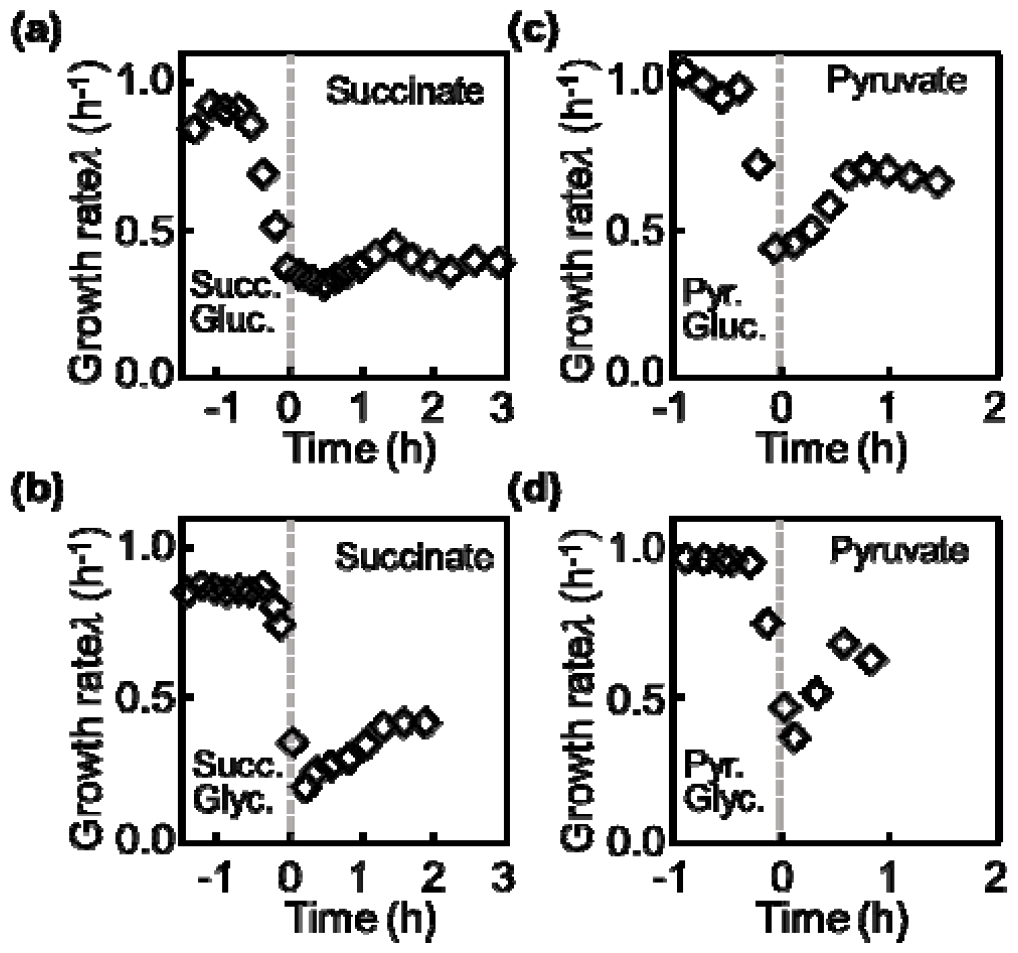
Overshoot of growth rate reported in other down-shift co-utilized carbon substrates. *E*.*coli* was grown on succinate (succ.) (a, b) or pyruvate (pyr.) (c, d) combined with either gluconate (gluc.) (a, c) or glycerol (glyc.) (b, d). At around t=0, gluconate or glycerol was depleted. Data was summarized from Erickson et al. [2].

**Fig. S10.**
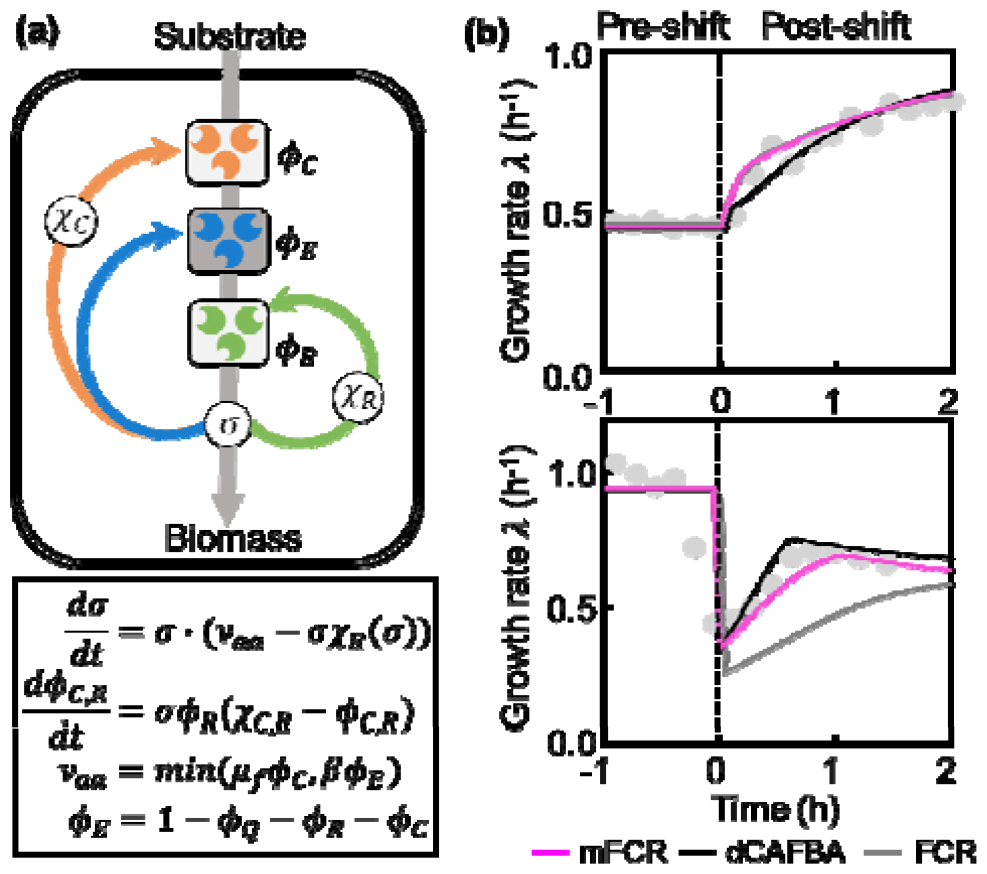
Model of the modified FCR. (a) Schematic view of the simplified model by integrating, which is a modification of the FCR model (mFCR). The carbon substrates are imported by carbon uptake proteins, which are then converted into central precursors by metabolic enzymes. The produced precursors are consumed by the ribosome for protein synthesis which determines the translational activity. Finally, the protein synthesis flux is allocated to coarse-grained sectors of catabolic (C), ribosomal (R) proteins via the regulatory functions and, respectively. (b) Comparisons of growth rate predicted by modified FCR, dCAFBA and FCR models in the up-shift (left) and down-shift (right). The magenta, black, grey curves are predictions of the mFCR, FCR and dCAFBA models, respectively. Grey dots indicate experimental data obtained from Erickson et al. [2]. For the up-shift, the predictions of mFCR (magenta curve) and FCR (grey curve) are overlapped. The simulated results of FCR in the down-shift did not match the experimental results due to the lack of consideration for nutrient depletion in the simulation. In the up-shift case, gluconate is added to the medium containing succinate. For the down-shift case, gluconate is depleted in the medium containing both pyruvate and gluconate.

**Fig. S11.**
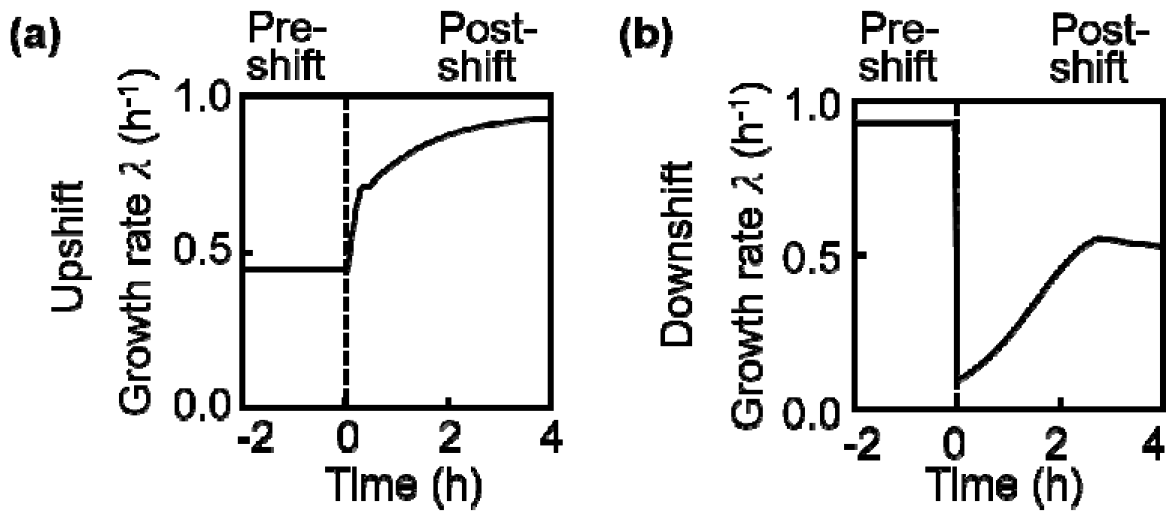
dCAFBA simulations of growth rate using the *E*.*coli i*ML1515 metabolic model during up-shift (a) and down-shift (b) with the carbon shift as figure 1b.

**Fig. S12.**
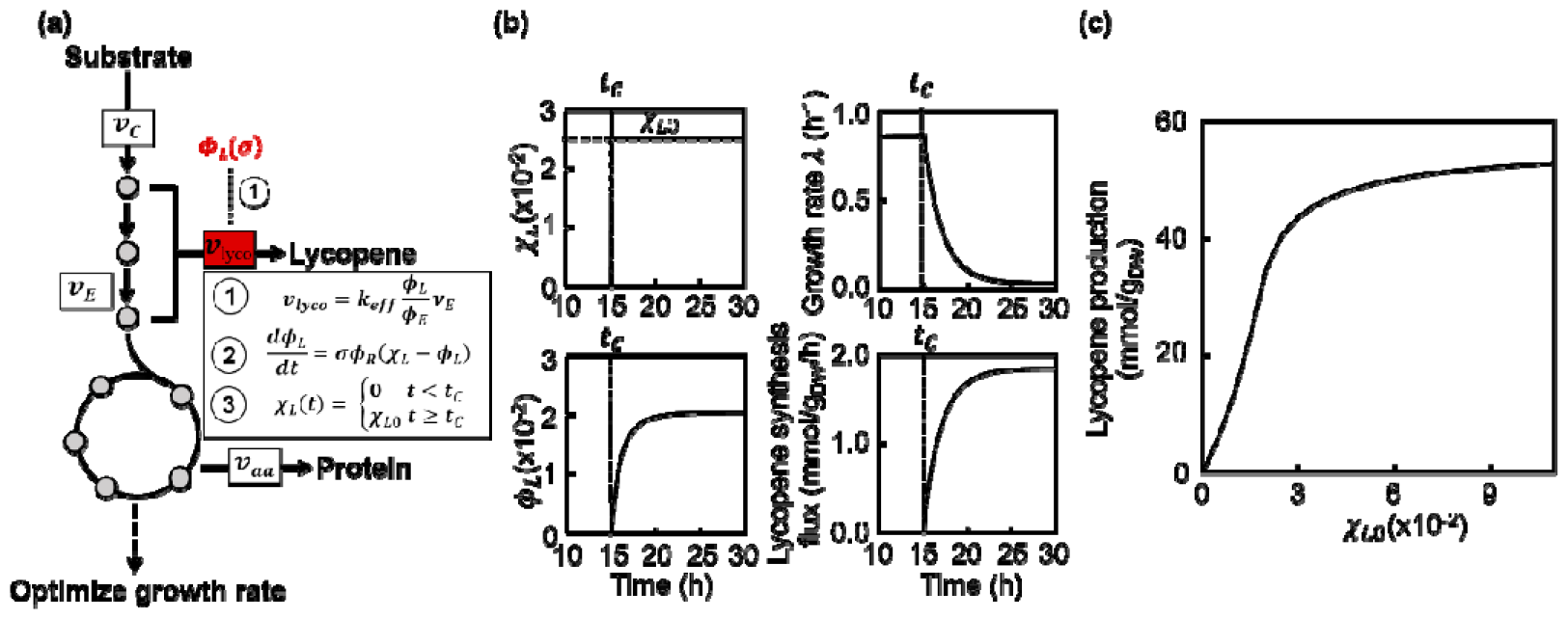
Applications of dCAFBA. (a) Schematic illustration of the dCAFBA model to simulate the growth dynamics perturbed by enhanced lycopene production in *E*.*coli*. (b) Regulation function of lycopene synthesis flux, protein fraction of lycopene synthesis, growth rate and lycopene synthesis upon inducing exogenous genes at with. (c) Changes of lycopene production accumulated in 24 h after induction under varying .

